# A spatial single-cell type multiplex map of human spermatogenesis

**DOI:** 10.1101/2024.10.21.619380

**Authors:** Feria Hikmet, Loren Méar, Jonas Gustavsson, Gisele Miranda, Cheng Zhang, Borbala Katona, Rutger Schutten, Per Adelsköld, Kalle Von Feilitzen, Mattias Forsberg, Jan-Bernd Stukenborg, Mathias Uhlén, Cecilia Lindskog

**Affiliations:** Department of Immunology, Genetics and Pathology, Cancer Precision Medicine Research Unit, Uppsala University, SE-75185 Uppsala, Sweden; Department of Organismal Biology, Nordfertil Research Lab Uppsala, Physiology and Environmental Toxicology, Uppsala University, SE-75185 Uppsala, Sweden; Department of Women’s and Children’s Health, Division for Neonatology, Obstetrics, Gynaecology and Reproductive Health, Karolinska Institute and Karolinska University Hospital, SE-14186 Stockholm, Sweden, Sweden; Science for Life Laboratory, Division of Computational Science and Technology, KTH Royal Institute of Technology; Roger Williams Institute of Liver Studies, School of Immunology & Microbial Sciences, Faculty of Life Sciences and Medicine, King’s College London, Foundation for Liver Research, and King’s College Hospital, London, United Kingdom; Science for Life Laboratory, School of Engineering Sciences in Chemistry, Biotechnology and Health, KTH Royal Institute of Technology, 17121 Stock-holm, Sweden; Department of Women’s and Children’s Health, Nordfertil Research Lab Stockholm, Childhood Cancer Research Unit, Karolinska Institutet, and Karolinska University, Stockholm, Sweden; Department of Neuroscience, Karolinska Institutet, Stockholm, Sweden

**Keywords:** spatial biology, antibody-based proteomics, single-cell profiling, testis, multiplex imaging, spermatogenesis

## Abstract

Understanding how transcriptional programs are translated into functional protein landscapes within intact tissues remains a major challenge in biology. While single-cell RNA sequencing (scRNA-seq) has enabled high-resolution identification of cellular states, corresponding protein expression within its spatial context remains poorly resolved at scale. Here, we present a scalable framework that integrates scRNA-seq with multiplex immunohistochemistry (mIHC) and automated image analysis to achieve spatially resolved, single-cell-level protein mapping across human spermatogenesis.

Using this approach, we quantified the expression of nearly 500 proteins across 12 discrete germ cell states within intact human testis tissue, generating a high-resolution spatiotemporal proteomic atlas. Comparative analysis of mRNA and protein expression revealed widespread temporal discordance, indicating that transcriptional state is not always predictive of protein abundance. Notably, we identify delayed translation of key regulators such as PIWIL4, whose protein expression peaks at later differentiation stages than its mRNA expression.

Together, our results establish a generalizable strategy for proteome-wide spatial mapping of single-cell states and demonstrate the importance of integrating protein-level measurements to resolve cellular identity and function in complex tissues.

## Background

Recent advances in single-cell RNA sequencing (scRNA-seq) have enabled unprecedented resolution in defining cellular identities across human tissues, resulting in multiple large-scale mapping efforts, including the Human Cell Atlas (HCA) ^1^, the Chan Zuckerberg Initiative (CZI) ^2^, the Human BioMolecular Atlas Program (HuB-MAP) ^3^, and the Human Cell Landscape ^4^. While transcriptomic analysis provides high sensitivity for detecting low-abundance genes, it is affected by technical limitations such as dropouts ^5^. In contrast, proteomic methods offer a broader dynamic range ^6^. Importantly, transcriptional profiles alone provide an incomplete view of cellular function, as protein abundance, localization, and timing are governed by complex post-transcriptional mechanisms ^7, 8^.

Bridging the gap between transcriptional state and functional protein output in a preserved spatial tissue context remains a central challenge in single-cell biology. Current approaches address this problem only partially. While scRNA-seq captures cellular heterogeneity at scale, it lacks spatial information and does not directly measure protein abundance. Spatial transcriptomics methods preserve tissue context but remain limited in sensitivity and do not resolve protein-level regulation. Conversely, antibody-based proteomics enables direct *in situ* measurements of proteins at a high spatial resolution, but traditional immunohistochemistry (IHC) is low-throughput and lacks the ability to quantitatively and systematically profile large numbers of proteins ^9^. In addition, identification of cell types relies on morphological features, and transition between molecularly defined single-cell states cannot always be resolved by histology alone ^10^. Finally, manual interpretation of images is inherently subjective and may introduce bias, further underscoring the need for more advanced imaging approaches ^11^.

The human testis provides an ideal system to address this challenge. Spermatogenesis is a highly ordered process accompanied by tightly regulated transcriptional and translational programs, during which the undifferentiated germ cells undergo meiosis, chromatin condensation, and extensive cellular remodeling ^12^. As differentiation progresses, sperm cells acquire specialized structures such as the flagellum and acrosome, enabling motility and fertilization ^13^. Notably, transcription is temporally restricted during key phases of meiosis and spermiogenesis, leading to the storage and delayed translation of mRNAs. This makes spermatogenesis particularly well-suited for investigating RNA-protein dynamics within a spatially organized tissue. In addition, the testis exhibits the highest number of tissue-specific genes among human tissues ^11, 14, 15^, many of which remain functionally uncharacterized ^16^. Since spermatogenesis is a continuous differentiation process, defining discrete germ cell types and states remains challenging. Traditional classification relies largely on morphology, such as nuclear condensation in spermatogonia or the shape of round and elongated spermatids. However, morphology alone cannot reliably resolve molecularly defined cell states, and as we have shown previously, manual image interpretation may introduce subjectivity and bias ^11, 15^. These limitations highlight the need for quantitative spatial methods that can link protein expression to precisely defined germ cell states.

Recent advances in multiplex protein imaging have enabled improved analysis of co-expression and tissue heterogeneity. Methods such as CODEX, Immuno-SABER, InSituPlex, and TOF-based approaches (including imaging mass cytometry and mass ion beam imaging) allow simultaneous detection of multiple markers. However, these approaches typically rely on predefined antibody panels and require complex antibody conjugation procedures, such as DNA barcoding. As a result, they are less suitable for flexible, large-scale mapping of protein expression across diverse cell states and are not readily applicable to proteome-wide analyses.

Here, we introduce a scalable framework for spatial single-cell proteomics that integrates scRNA-seq-defined cell states with multiplex IHC (mIHC) and automated image analysis. By combining fixed antibody panels for cell-state identification with iterative staining of candidate proteins, we enable quantitative, single-cell-resolved mapping of protein expression within intact tissue. Applying this approach to nearly 500 proteins in human testis, we generate a comprehensive spatial proteomic atlas spanning 12 biologically defined germ cell states across spermatogenesis and systematically compare mRNA and protein expression dynamics. All spatial proteomic data are made publicly available through the Human Protein Atlas (HPA) (www.proteinatlas.org) ^14, 17^. Our analysis reveals widespread discordance between transcript and protein expression across cell states, highlighting the importance of post-transcriptional regulation in defining cellular identity. Beyond providing a detailed map of human spermatogenesis, this work establishes a broadly applicable strategy for integrating transcriptomic and proteomic data to resolve cellular function in complex tissues.

## Results

### A spatial single-cell framework for mapping the human testis proteome

To map protein expression across molecularly defined germ cell states in intact tissue, we developed an integrated workflow combining antibody-based protein profiling, scRNA-seq, mIHC, and automated image analysis (Figure 1). This framework enables quantitative, spatially resolved protein mapping at single-cell resolution across the complex architecture of the human testis. Spermatogenesis occurs within a highly dynamic and heterogeneous tissue environment, in which spermatogonial stem cells differentiate into mature sperm through tightly coordinated developmental stages supported by surrounding niche cells (Figure 1A). To capture this complexity, we first generated a high-resolution, image-based atlas of the testis proteome by integrating IHC data with scRNA-seq defined cell states (Figure 1B). This multimodal reference was used to design multiplex antibody panels for spatially resolving germ cell states. Using these panels, we performed iterative mIHC to map protein expression of candidate targets across defined cell states within intact tissue sections (Figure 1C). To enable quantitative analysis, we developed an automated image analysis pipeline that performs cell segmentation, phenotyping, and protein expression quantification at single-cell resolution (Figure 1D). This approach provides a scalable platform for systematic interrogation of protein expression patterns and functional relationships across spermatogenesis (Figure 1E).

**Fig 1.**
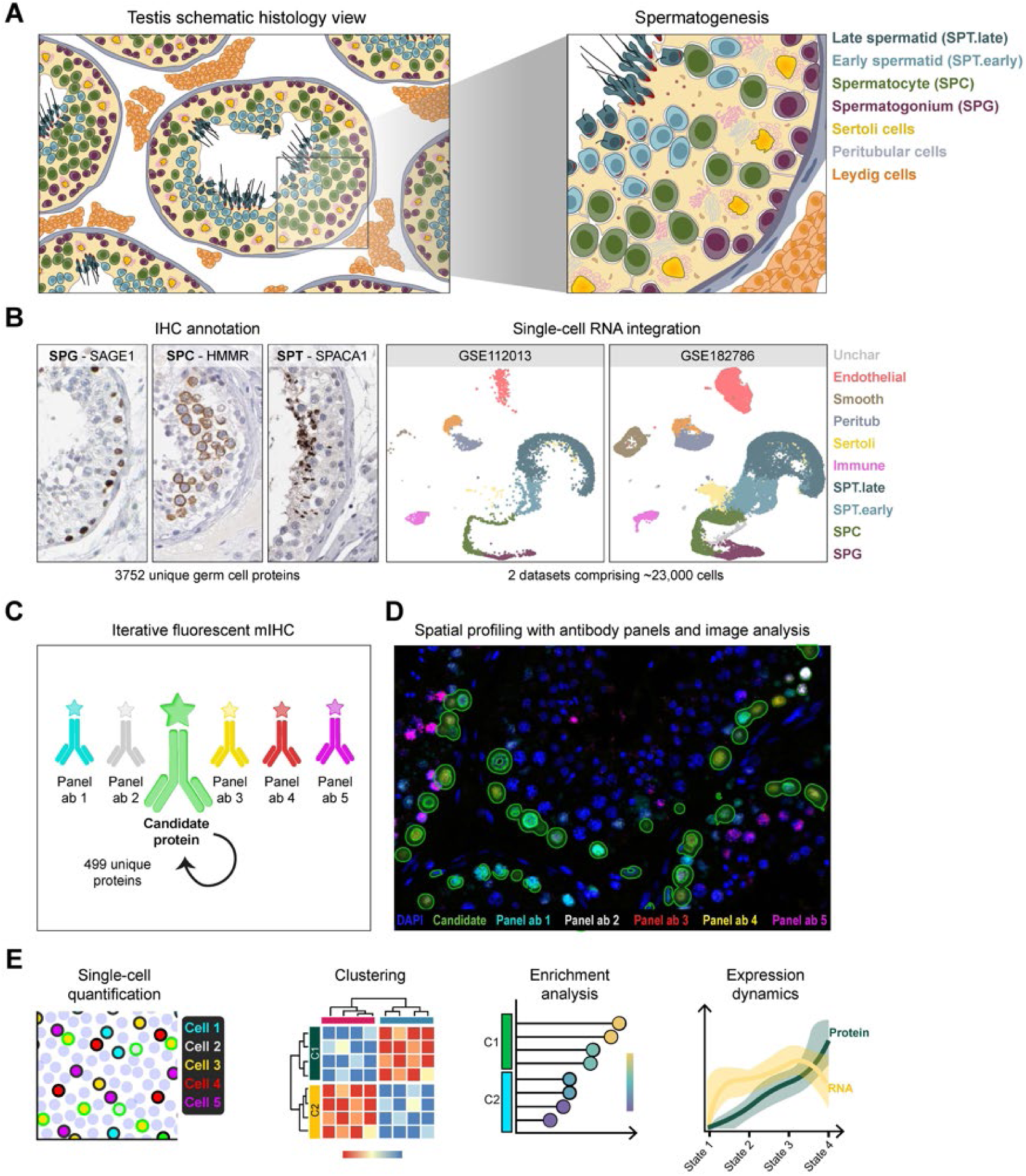
Overview of the study workflow. (**A**) Schematic representation of the complex testis microstructure, illustrating germ cells within the seminiferous tubules. (**B**) Integration of immunohistochemistry (IHC)-based annotation and single-cell RNA sequencing (scRNA-seq) datasets to construct multiplex IHC (mIHC) antibody panels for spatial characterization of germ cell proteins. (**C**) Iterative staining of candidate proteins together with fixed panel markers for cell state identification; the candidate protein is shown in green (see Methods for details). (**D**) Automated image analysis pipeline for cell segmentation and phenotyping, enabling quantitative assessment of candidate protein expression at single-cell resolution (green overlays). (**E**) Downstream analyses based on high-resolution single-cell data, including protein quantification across cell states and functional characterization.

### Integrated antibody profiling and scRNA-seq define germ cell states

To construct a high-resolution spatial proteomics reference of the human testis, we leveraged an IHC-based dataset comprising 4,431 proteins analyzed using validated antibodies. Of these, 3,752 unique proteins showed detectable expression in at least one germ cell type and were retained for downstream analysis. For proteins assessed with multiple antibodies (n = 1,124), the antibody with the highest signal-to-noise ratio was selected (Figure 2A). This curated set formed the basis for selecting both panel markers and candidate proteins.

**Fig 2.**
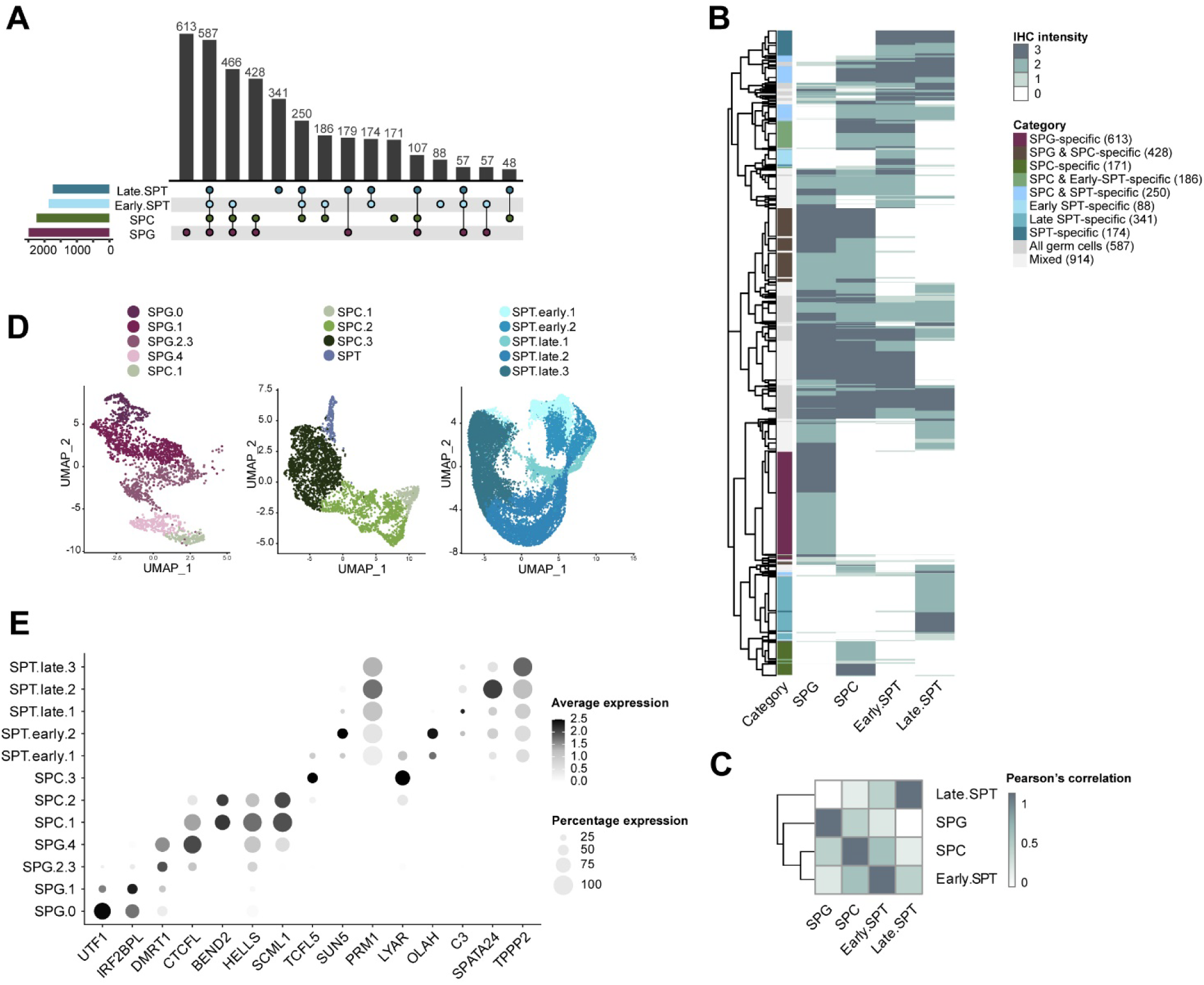
The tissue-based testis proteome defined by antibody profiling and single-cell transcriptomics. (**A**) IHC annotation intensity values were binarized (negative = 0; weak, moderate, and strong = 1) and visualized as an UpSet plot. Horizontal bars indicate the number of proteins detected in each germ cell type, while vertical bars represent the frequency of specific combinations of protein expression across cell types. Total counts are shown above each bar. (**B**) Heatmap with hierarchical clustering showing IHC staining intensity (negative, weak, moderate, strong) across germ cell types (columns) for 3,752 proteins (rows). Proteins are annotated by cell type specificity groups, with counts indicated in parentheses. (**C**) Spearman correlation matrix based on the data shown in (B), with color indicating correlation coefficients between germ cell types. **(D)** UMAP projections of scRNA-seq sub-clustering for spermatogonia (left), spermatocytes (middle), and spermatids (right). **(E)** Dot plot showing selected marker genes used for mIHC panel design. The x-axis represents genes, and the y-axis shows their mean expression and distribution across defined sub-clusters. Genes are ordered according to their position in the schematic panel design: UTF1, IRF2BPL, DMRT1, CTCFL, and BEND2 for the spermatogonia panel (SPG.0 to SPC.1); HELLS, SCML1, TCFL5, SUN5, and PRM1 for the spermatocyte panel (SPC.1 to SPT.late); and LYAR, OLAH, C3, SPATA24, and TPPP2 for the spermatid panel (SPT.early.1 to SPT.late.3).

Analysis of staining patterns revealed both cell type-specific and broadly expressed proteins across spermatogenesis. Distinct subsets displayed strong specificity to spermatogonia, spermatocytes, or spermatids, whereas others exhibited more gradual, stage-dependent expression patterns (Figure 2B). Globally, protein expression profiles were highly correlated across germ cell types, with the strongest similarity observed between spermatocytes and early spermatids (Figure 2C), reflecting shared transcriptional and functional programs.

To define the cellular states underlying these expression patterns, we integrated scRNA-seq data from two independent datasets ^18, 19^. After quality control, 37,022 cells were retained and grouped into major cell types, including spermatogonia (SPG), spermatocytes (SPC), spermatids (SPT), and somatic niche cells (Supplementary Figure 1). Clustering analysis identified 26 transcriptionally distinct populations, and pseudotime trajectory analysis confirmed a continuous progression from undifferentiated to differentiated germ cell states.

Focusing on germ cells, sub-clustering identified 12 SPG, 14 SPC, and 16 SPT transcriptional sub-clusters. These were consolidated into 12 biologically interpretable germ cell states for spatial protein mapping based on marker expression and differentiation stage. (Supplementary Figure 2). Spermatogonia were classified into four main states (SPG.0, SPG.1, SPG.2-3, and SPG.4), including a transitional population expressing spermatocyte markers (SPC.1). Spermatocytes were resolved into three principal meiotic stages (SPC.1-3), while spermatids comprised early (SPT.early.1-2) and late (SPT.late.1-3) differentiation states associated with progressive acquisition of sperm-specific features (Figure 2D and Supplementary Figure 2).

To enable spatial mapping of these cell states, we identified candidate marker genes using differential expression analysis (fold-change > 1.5). Among the proteins with available antibodies in the HPA, 969 differentially expressed genes (DEGs) were associated with spermatogonial states, 2,093 with spermatocytes, and 1,332 with spermatids. From these, representative markers were selected to construct three mIHC panels for SPG, SPC, and SPT states (Figure 2E). Where possible, well-established markers (e.g., UTF1 for SPG.0 and DMRT1 for SPG.2-3) were prioritized. For less well-characterized states, candidates were selected based on staining quality and spatial specificity in IHC data, ensuring compatibility with downstream automated image analysis.

### Multiplex imaging maps nearly 500 proteins across spermatogenesis

To enable spatial mapping of germ cell states, we developed mIHC antibody panels that robustly and specifically label distinct spermatogenic populations. Panel markers were selected and optimized to ensure clear discrimination of cell states corresponding to scRNA-seq-defined sub-clusters, with minimal overlap between markers within each panel (Figure 3A and Supplementary Figure 3). To further refine state boundaries, exclusion markers were incorporated, including a preleptotene spermatocyte marker in the SPG panel and early and late spermatid markers in the SPC panel.

**Fig. 3.**
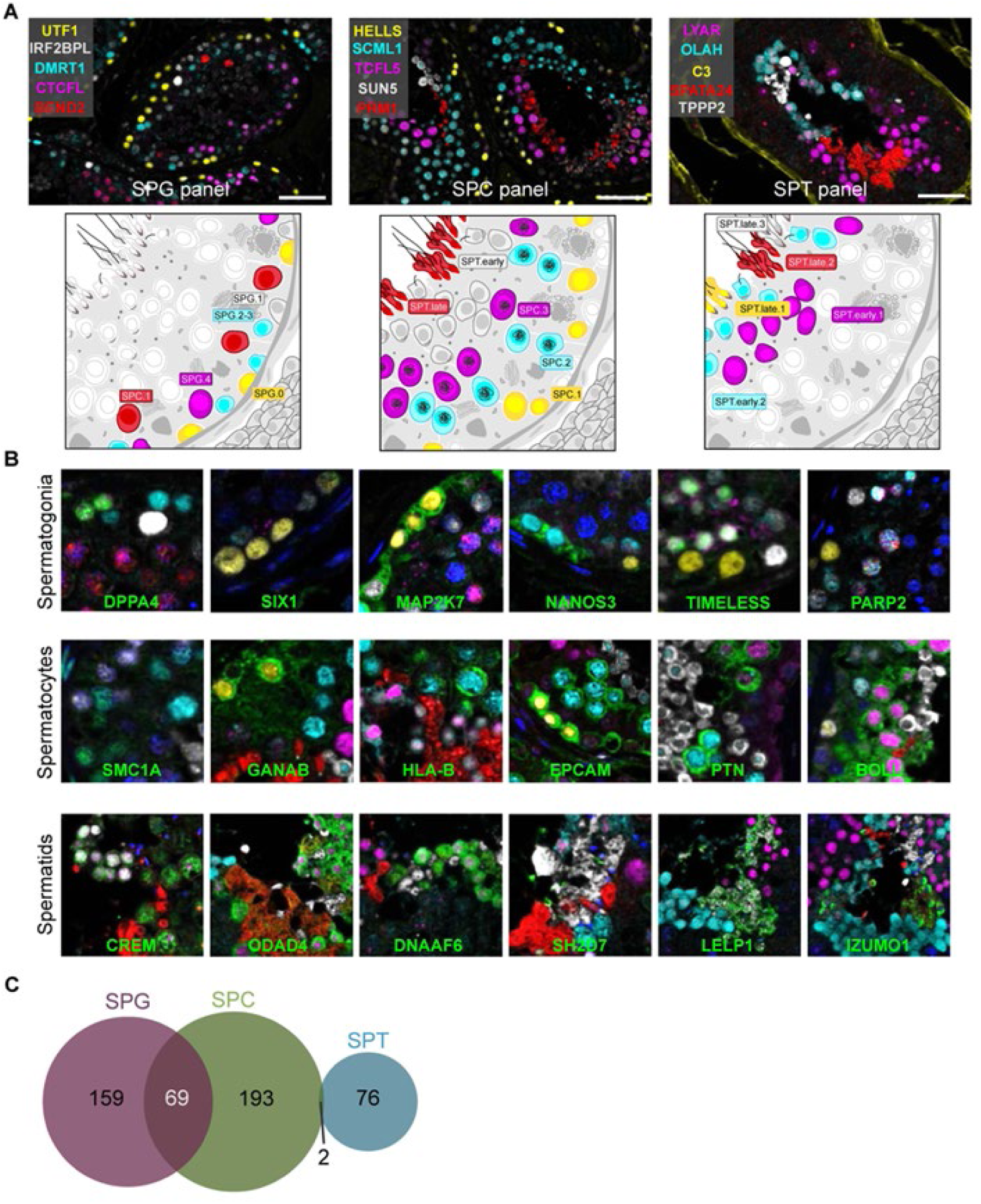
Deep spermatogenic profiling using high-throughput multiplex immunohistochemistry. (**A**) Representative images of each mIHC panel in normal human seminiferous tubules, with schematic illustrations indicating the relative abundance and spatial organization of the targeted germ cell states. (**B**) Representative examples of candidate proteins stained using the three antibody panels. Candidate proteins are shown in green, with protein names indicated in each image. (**C**) Venn diagram showing the number of unique candidate proteins analyzed with each panel.

Using these panels, we performed iterative mIHC to map protein expression across defined germ cell states. In each staining cycle, a single candidate protein was combined with a fixed panel, enabling assignment of its expression to specific cell states based on co-localization with panel markers (Figure 3B). This strategy allowed systematic profiling of protein localization across spermatogenesis at single-cell resolution. In total, 499 candidate proteins were selected for analysis based on prior IHC-defined expression patterns (Figure 1B-C; Supplementary Table 3). Panel assignment was guided by expected cell type specificity, ensuring that proteins were analyzed in the most relevant cellular context (Supplementary Figure 4). To capture transitional expression dynamics, 69 proteins were analyzed across multiple panels where cell states overlapped (e.g., SPC.1), enabling precise determination of expression onset and termination across differentiation.

Overall, 228 proteins were mapped using the SPG panel, 264 using the SPC panel, and 78 using the SPT panel (Figure 3C). Together, this dataset represents a large-scale spatial proteomic resource spanning nearly 500 proteins across defined stages of human spermatogenesis.

### Automated image analysis enables quantitative single-cell protein profiling

mIHC generated high-resolution, multi-channel fluorescence images for each of the 499 candidate proteins. To enable quantitative analysis across these datasets, we developed a custom image analysis pipeline that performs cell segmentation, phenotyping, and protein expression quantification at single-cell resolution (Figure 4A). Using this framework, individual cells were first identified and segmented, followed by classification into defined germ cell states based on panel marker expression. This enabled consistent cell state annotation across all three antibody panels, with the relative distribution of cell states shown in Figure 4B. Protein expression was then quantified within each phenotyped cell using a machine learning-based approach. To distinguish true signal from background, a pixel-level classifier was trained on a subset of annotated images and subsequently applied across the full dataset to generate masks of positive protein expression (Supplementary Figure 5). This approach enabled robust removal of non-specific signal and standardized quantification of protein intensity across all samples.

**Fig. 4.**
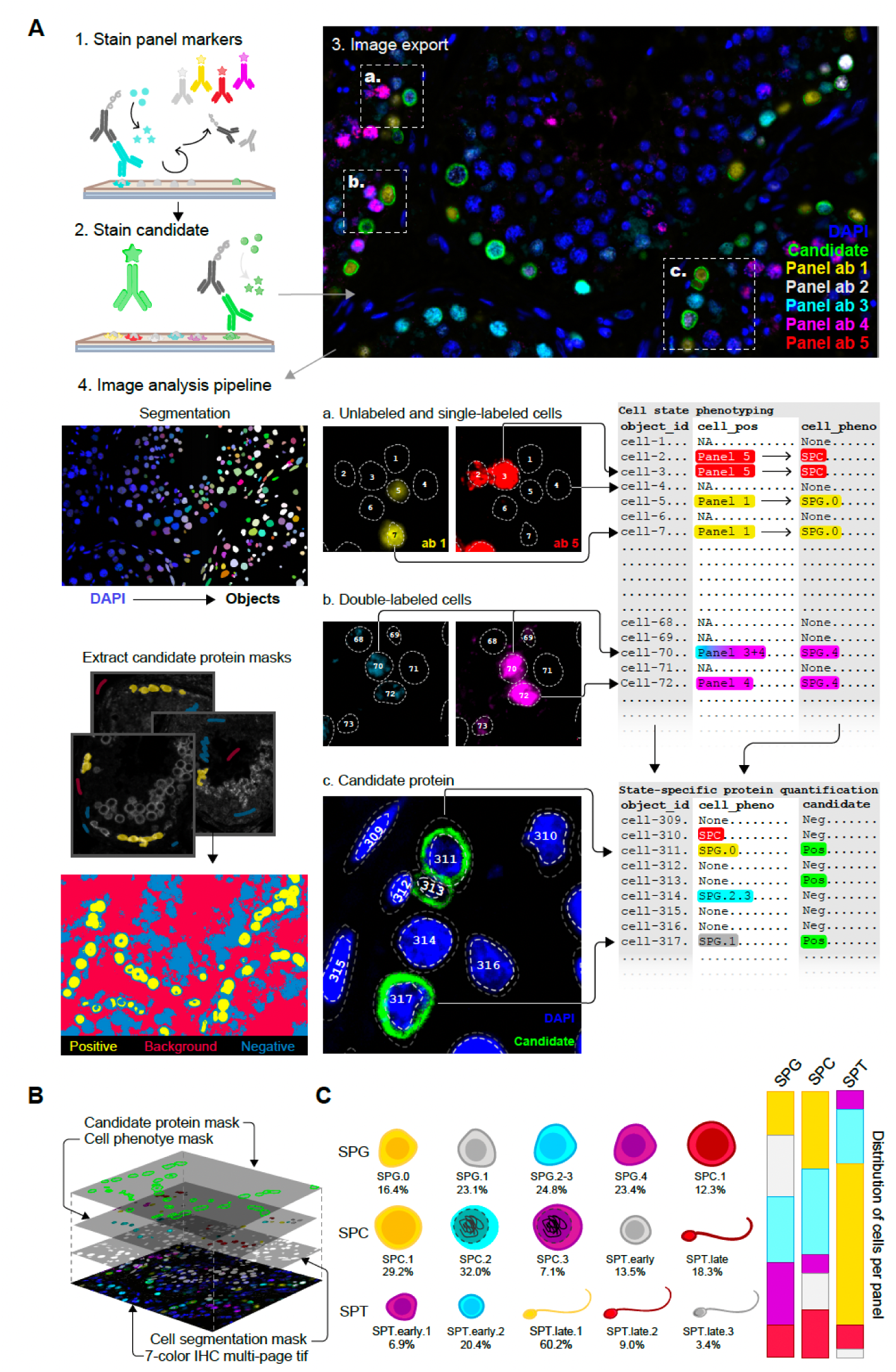
High-resolution quantification and single-cell characterization by bioimage analysis. (**A**) Workflow for image processing and single-cell quantification. Multiplex-stained testis sections were imaged by fluorescence microscopy and spectrally unmixed using panel-specific algorithms. An example image shows LBR (green) stained with the SPG panel. Nuclei were segmented based on the DAPI channel to define individual cells. Candidate protein signals were classified using a pixel-based machine learning model to identify positive signal, which was used to generate cell-level masks. In parallel, panel marker signals were thresholded to assign cell identities, including single- and double-positive populations. These data were integrated to generate a table linking protein expression to cell state for each cell. (**B**) Overview of the different masks generated from each multiplex image, including nuclear segmentation, marker-based cell phenotyping, and candidate protein signal detection. (**C**) Schematic representation of cell states identified within each panel, shown with pseudo-colors, and the corresponding distribution of cell states across all analyzed proteins.

Together, this pipeline provides a scalable and reproducible framework for extracting quantitative, single-cell-resolved protein expression data from multiplex imaging experiments. The resulting dataset includes cell counts and mean expression intensities for each protein across all cell states (Supplementary Tables 8-9).

### Protein expression programs define functional stages of spermatogenesis

To define protein expression programs across spermatogenesis, we performed hierarchical clustering of candidate proteins based on their cell state-specific expression profiles. Functional annotation of the resulting clusters using Gene Ontology (GO) enrichment analysis revealed distinct stage-associated protein programs spanning germ cell differentiation.

Among the 228 proteins analyzed with the SPG panel, five major clusters were identified based on their cell state-specific expression patterns (Figure 5A, Supplementary Table 4). Early spermatogonial states were represented by Cluster 2, the largest group (81 proteins), which showed strong enrichment in SPG.0 and SPG.1 cells and was associated with transcriptional regulation. Notably, these proteins showed little to no expression in SPC.1, indicating restriction to undifferentiated spermatogonia. Intermediate and late spermatogonial states were captured by two distinct clusters. Cluster 1 (52 proteins) was primarily associated with SPG.2–3 cells and enriched for lamellipodium-related functions. Cluster 4 (53 proteins) showed specific expression in SPG.4 cells and was linked to cell cycle regulation, consistent with progression toward spermatocyte entry. A smaller group of proteins in Cluster 5 (23 proteins) displayed expression in both SPG.0 and SPC.1 cells, suggesting a transitional program between spermatogonial and early spermatocyte states, and was also enriched for transcription-related functions. Finally, Cluster 3 (19 proteins) exhibited a mixed expression pattern across multiple states without a clearly dominant functional annotation.

**Fig. 5.**
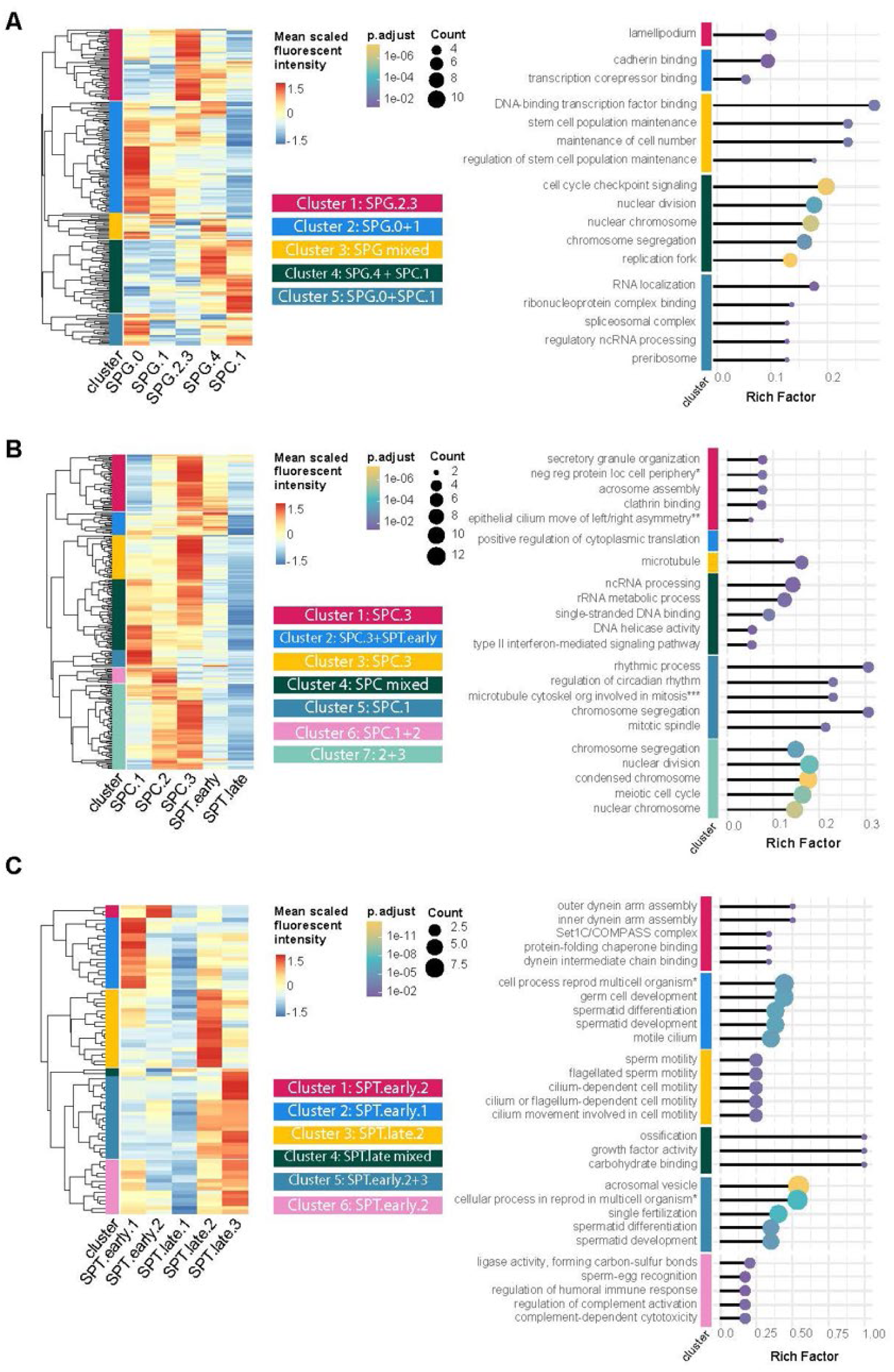
Hierarchical clustering and functional characterization of protein expression across germ cell states. Protein expression profiles are shown separately for **(A)** spermatogonia (SPG), **(B)** spermatocytes (SPC), and **(C)** spermatids (SPT). Heatmaps display row-wise z-score–normalized protein intensity values. Hierarchical clustering grouped proteins based on their cell state-specific expression patterns, and clusters were defined using cut-tree segmentation and numbered from top to bottom. Functional annotation of each cluster was performed using Gene Ontology (GO) enrichment analysis. Selected GO terms marked with an asterisk (*) are abbreviated for clarity.

Next, proteins analyzed with the SPC panel were subjected to hierarchical clustering (Figure 5B, Supplementary Table 4), revealing both cell state-specific and broadly expressed patterns. Overall, protein expression was strongly enriched in SPC.3 cells, which dominated several clusters. Clusters 1 (48 proteins) and 3 (37 proteins) were both characterized by high expression in SPC.3 cells, although Cluster 3 showed relatively lower expression in SPC.2, while Cluster 1 displayed reduced expression in SPT.early cells. In addition, Cluster 2 (19 proteins) included proteins expressed in both SPC.3 and SPT.early cells, indicating a transitional expression pattern between spermatocyte and early spermatid states. Other clusters were associated with earlier spermatocyte stages. Cluster 7, the largest group (73 proteins), was predominantly expressed in SPC.2 cells. Similarly, Cluster 6 (13 proteins) showed strong specificity for SPC.2, whereas Cluster 5 (14 proteins) was primarily restricted to SPC.1 cells. Finally, Cluster 4 (60 proteins) exhibited a broader expression pattern across multiple spermatocyte states without clear restriction to a single stage.

Functional annotation of the SPC clusters by GO enrichment analysis indicated that these proteins are primarily involved in RNA processing, chromatin organization, and structural chromosomal changes associated with meiosis (Figure 5B).

Proteins analyzed with the SPT panel were grouped into six clusters reflecting distinct stages of spermiogenesis (Figure 5C, Supplementary Table 4). Early spermatid states were represented by Cluster 1 (3 proteins) and Cluster 2 (18 proteins), which were associated with SPT.early.1 and SPT.early.2 cells. These clusters capture proteins involved in the initial phases of haploid cell differentiation. Intermediate stages were reflected by Cluster 3 (20 proteins), which showed predominant expression in SPT.late.2 cells, indicating progression toward structural maturation. Late spermiogenesis was dominated by Clusters 5 (21 proteins) and 6 (14 proteins), which exhibited strong expression in SPT.late.2 and SPT.late.3 cells. These clusters represent proteins involved in the final stages of sperm development. Cluster 4 (2 proteins) displayed a similar pattern but with more pronounced expression in SPT.late.1 cells, suggesting an intermediate late-stage profile.

Functional annotation of the SPT clusters indicated enrichment for processes related to sperm flagellum assembly, motility, fertilization, and acrosome formation (Figure 5C).

To illustrate cluster-specific expression patterns, representative proteins from each SPG, SPC, and SPT cluster are shown together with quantitative profiles in Figure 6.

**Fig. 6.**
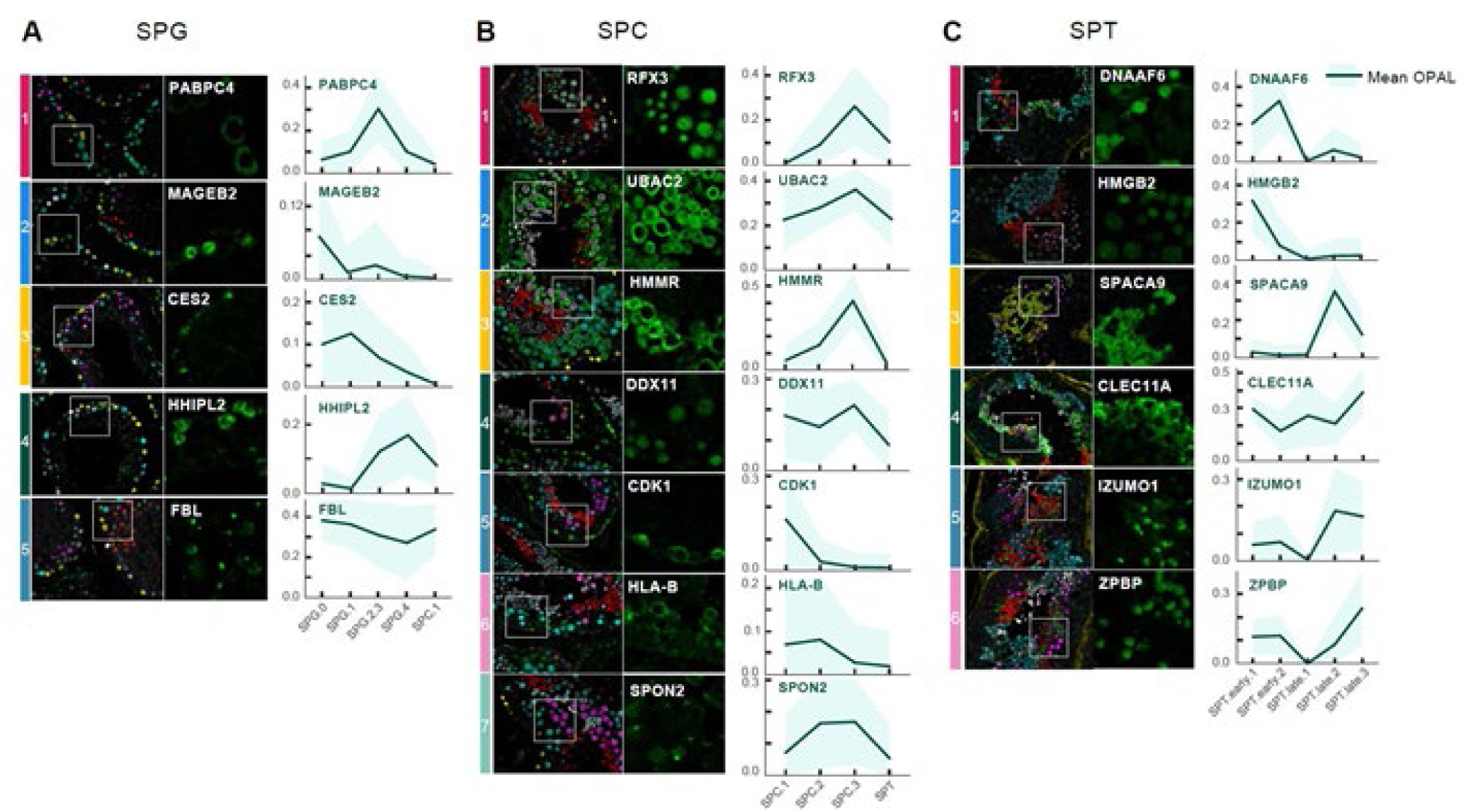
Representative protein expression patterns across spermatogenic states. Examples of proteins from each expression cluster are shown for **(A)** spermatogonia (SPG), **(B)** spermatocytes (SPC), and **(C)** spermatids (SPT). For each protein, multiplex images (left) display all panel markers, while corresponding insets (right) show the candidate protein signal (green). Line plots illustrate normalized protein expression across cell states based on image analysis quantification. Cluster assignments correspond to the hierarchical clustering results shown in Figure 5.

Within the SPG panel, PABPC4 (Cluster 1), a regulator of mRNA metabolism ^20^, was enriched in SPG.2-3 cells. MAGEB2 (Cluster 2), a cancer-testis antigen linked to transcriptional regulation ^21^, and CES2 (Cluster 3), involved in cholesterol metabolism ^22^, were primarily detected in early spermatogonia (SPG.0-1). HHIPL2 (Cluster 4) showed specific expression in SPG.4, whereas FBL (Cluster 5), a nucleolar protein involved in rRNA processing ^23^, displayed broader expression across SPG states and SPC.1.

A similar analysis of SPC clusters revealed distinct stage-associated patterns. RFX3 (Cluster 1), a regulator of ciliogenesis ^24^, was enriched in SPC.3 and early spermatids. UBAC2 (Cluster 2), associated with WNT signaling ^25^, exhibited broader expression across spermatocyte and spermatid states. HMMR (Cluster 3), involved in cell motility ^26^, showed specificity for SPC.3, while DDX11 (Cluster 4), required for chromosome cohesion ^27^, was most prominent in early spermatocytes. CDK1 (Cluster 5), a key regulator of the G2–M transition ^28^, was restricted to SPC.1, and HLA-B (Cluster 6), involved in antigen presentation ^29^, was enriched in SPC.2. SPON2 (Cluster 7), associated with cell adhesion ^30^, was detected across multiple states, with higher expression in SPC.2-3.

In SPT, early and late stages were similarly distinguished by cluster-specific protein expression. DNAAF6 (Cluster 1), required for flagellar motility ^31^, and HMGB2 (Cluster 2), involved in DNA remodeling and repair ^32^, were associated with early spermatid states. SPACA9 (Cluster 3), a dynein-associated microtubule protein ^33^, was enriched in SPT.late.2. CLEC11A (Cluster 4), linked to cell differentiation ^34^, showed broader expression with higher levels in later spermatid stages. IZUMO1 and ZPBP (Clusters 5-6), both essential for sperm–egg interaction and acrosome function ^35, 36^, were restricted to late spermatids (SPT.late.2-3).

### RNA-protein comparisons reveal temporal uncoupling during germ cell differentiation

Spermatogenesis is characterized by defined periods of transcriptional activity and inactivity, resulting in temporally separated waves of mRNA production ^37, 38^. We next used the spatial proteomic atlas to assess whether transcript abundance predicts protein levels across germ cell differentiation states (Figure 7A).

**Fig. 7.**
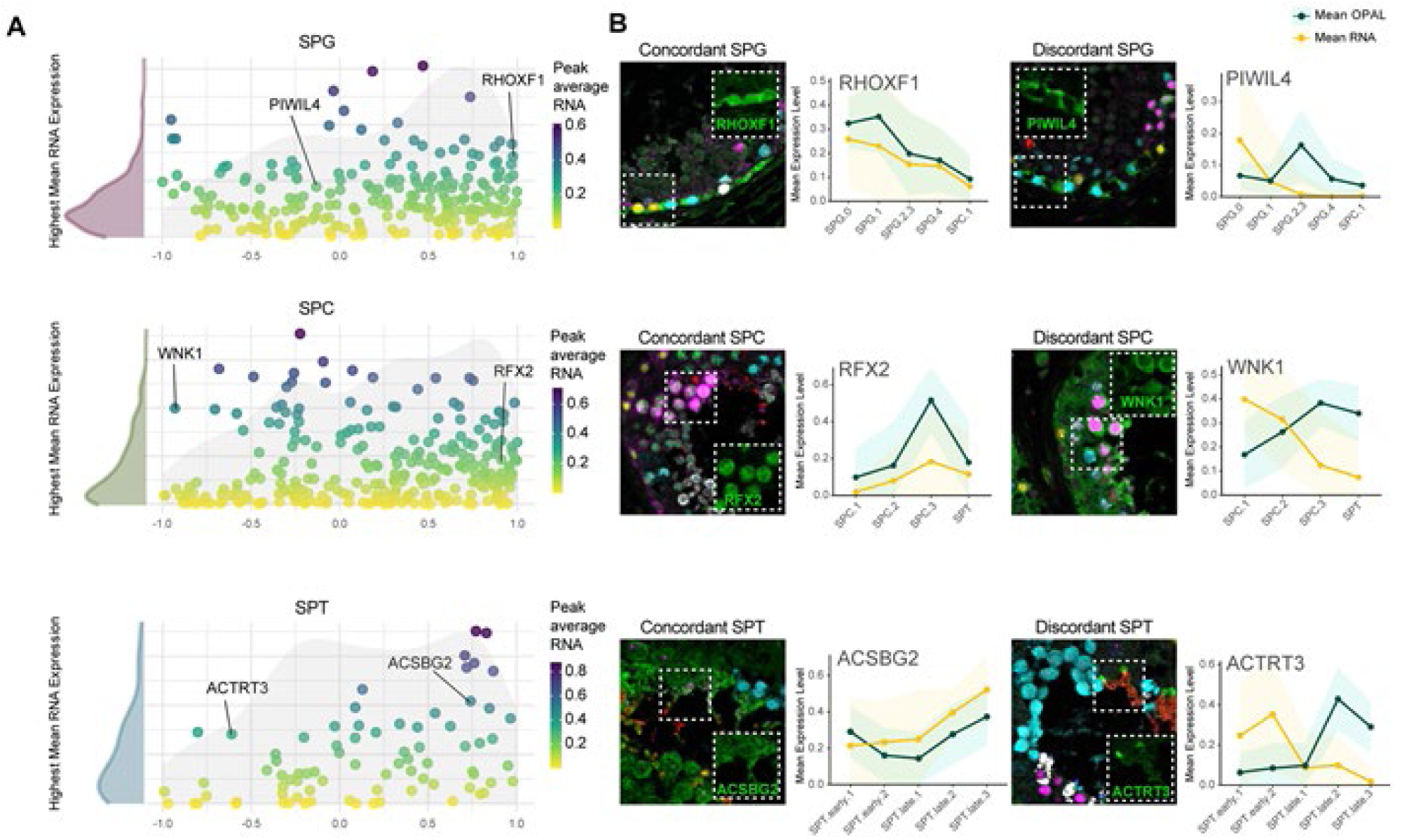
Spatiotemporal dynamics of mRNA and protein expression during spermatogenesis. (**A**) Pearson correlation between mRNA and protein levels at the single-cell level for each panel. Each dot represents a protein, with correlation coefficients (−1 to 1) calculated from scaled mRNA and protein expression across differentiation from least to most mature cell states within SPG, SPC, and SPT panels (y-axis). The x-axis indicates the cell state with the highest mean mRNA expression based on normalized scRNA-seq data. Density plots show the distribution of correlation coefficients (grey) and peak mRNA expression states (color). (**B**) Representative examples of concordant and discordant expression patterns. Multiplex images show candidate protein staining (green), and line plots display mean scaled mRNA (yellow) and protein (green) expression across cell states. Shaded areas indicate standard error.

Overall, most proteins showed positive RNA-protein correlations, indicating that mRNA is often translated within the same cell state in which it is transcribed. For example, the spermatogonial marker RHOXF1 exhibited concordant expression (Figure 7B), with both mRNA and protein peaking in the SPG.0 state. Similarly, the transcription factor RFX2 was enriched in SPC.3 at both the mRNA and protein levels, and ACSBG2 showed a consistent increase from SPT.early.1 to SPT.late.3 in both modalities (Figure 7B).

In contrast, several genes displayed discordant RNA-protein dynamics, where mRNA expression preceded protein accumulation. Notably, PIWIL4, previously described as a SPG.0 marker ^18^, showed strong mRNA enrichment in SPG.0, whereas the protein was predominantly detected in SPG.2-3 and SPG.4 (Figure 7A). A similar pattern was observed for WNK1, with transcript levels highest in SPC.1 and decreasing toward the spermatid stage, while protein expression peaked in SPC.3 (Figure 7B). Likewise, ACTRT3 transcripts were detected in early spermatids (SPT.early.1-2), whereas the protein accumulated at later stages, primarily in SPT.late.2-3 (Figure 7B). Finally, two proteins, VSIG4 (SPG panel) and R3HDML (SPC panel), could not be evaluated due to insufficient variability in the scRNA-seq data (Supplementary Table 9).

### RNAscope validates delayed PIWIL4 protein expression

To validate the observed RNA-protein relationships at single-cell resolution, we performed RNAscope combined with immunofluorescent staining for UTF1, PIWIL4, and RHOXF1 in human testis tissue sections (Figure 8A).

**Fig. 8.**
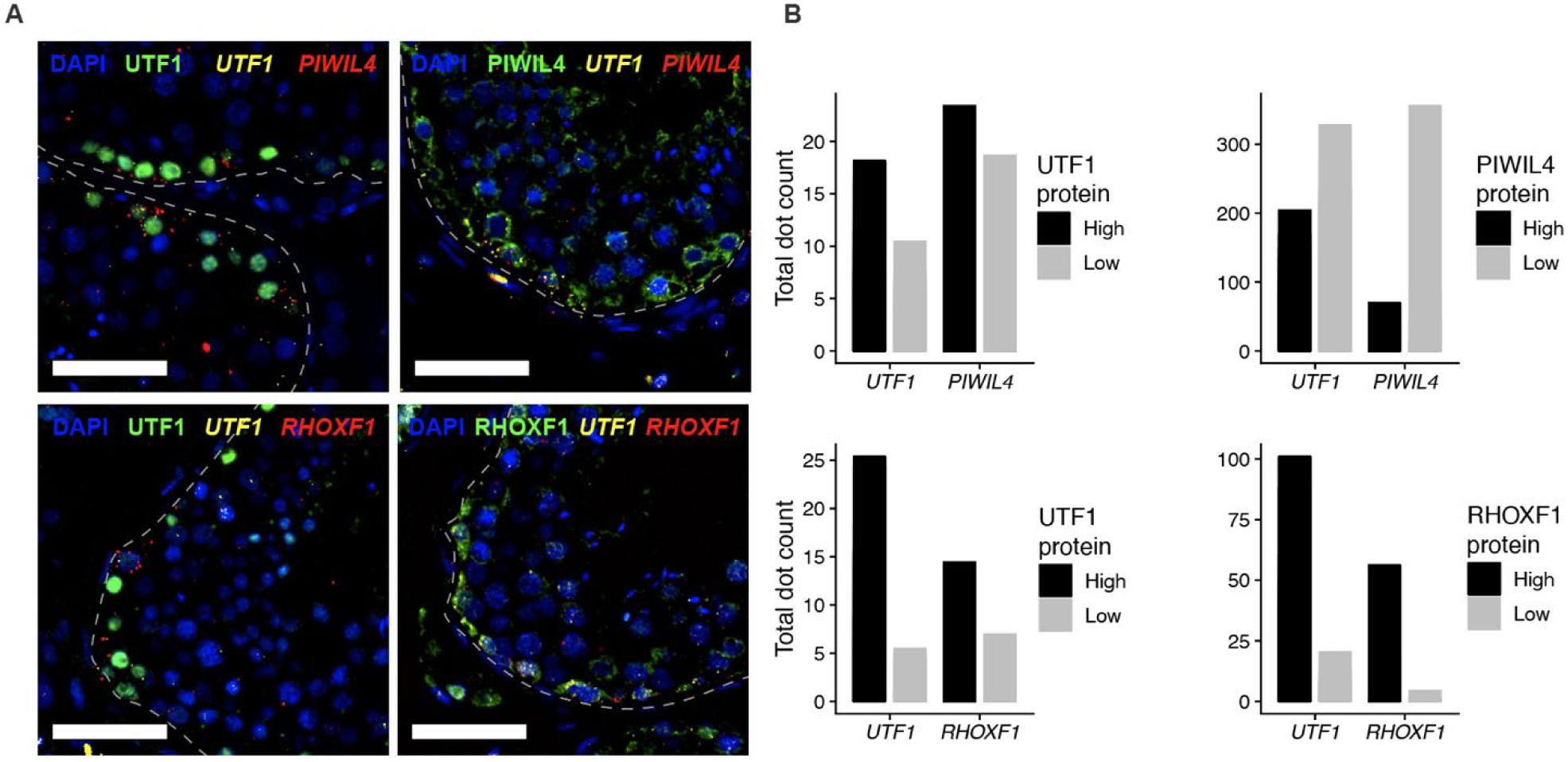
Validation of RNA-to-protein expression dynamics of PIWIL4 and RHOXF1 in SPGs using RNAscope. (**A**) Representative fluorescent images of RNA-protein co-detection showing *UTF1, PIWIL4* and *RHOXF1* RNA (yellow and red) and protein expression (green) in seminiferous tubules of the testis. Scale bar, 50µm. (**B**) Quantification of RNAscope signal (puncta per cell) in protein-high and protein-low cell populations.

Quantification of RNAscope signals showed that PIWIL4 and UTF1 transcripts were enriched in UTF1 protein-high spermatogonia, whereas both transcripts were reduced in PIWIL4 protein-high cells (Figure 8B, top). These results indicate that PIWIL4 mRNA is expressed in early spermatogonia, while protein accumulation occurs in more differentiated states, consistent with delayed translation. In contrast, RHOXF1 RNA signals were primarily detected in RHOXF1 protein-high spermatogonia (Figure 8B, bottom), supporting a concordant RNA-protein expression pattern.

## Discussion

Our integrated spatial proteomic and transcriptomic analysis reveals widespread temporal uncoupling between mRNA and protein expression during human spermatogenesis. These findings demonstrate that transcriptional state alone is not sufficient to predict protein abundance during germ cell differentiation. More broadly, linking transcriptional identity to functional protein output within intact tissues remains a central challenge in single-cell biology. While scRNA-seq has enabled high-resolution mapping of cellular states across human tissues ^1, 17^, corresponding protein-level information at comparable spatial and single-cell resolution has been lacking. By integrating transcriptomics with spatial proteomics, our study provides a scalable framework to systematically resolve RNA-protein relationships within their native tissue context.

Spermatogenesis represents a uniquely suitable system for studying RNA-protein dynamics, as it is governed by tightly coordinated transcriptional and translational programs. Transcription occurs in two major waves before and after meiosis, with the latter associated with storage of mRNAs for delayed translation. However, it has remained unclear which transcripts are translated immediately, and which are temporally uncoupled from protein production. Our analysis provides direct evidence that such temporal uncoupling is widespread across germ cell states, reinforcing the central role of post-transcriptional regulation in shaping the proteome during differentiation.

By combining scRNA-seq-defined cell states with mIHC and automated image analysis, we established a scalable framework for spatial single-cell proteomics. This approach enables quantitative mapping of nearly 500 proteins across 12 discrete germ cell states within intact tissue, thereby extending previous transcriptomic atlases to the protein level. Importantly, spatial analysis preserves tissue architecture and cellular context, avoiding biases introduced by tissue dissociation and enabling direct interpretation of protein expression in relation to differentiation stage and microenvironment. Our results reveal that protein expression is organized into stage-specific functional programs that reflect the biological processes underlying spermatogenesis. Early germ cell states are enriched for proteins associated with transcriptional regulation, whereas later stages are dominated by factors involved in chromatin remodeling, structural maturation, and sperm function. Many of the identified proteins have not previously been characterized in a spatial, cell state-resolved context, providing new insights into their potential roles in testicular biology.

A key observation is the frequent discordance between mRNA and protein expression across differentiation states. PIWIL4, previously regarded alongside UTF1 as a marker of naïve spermatogonia ^19, 39^, represents a prominent example. While PIWIL4 transcripts are enriched in early spermatogonia (SPG.0), we show that the protein accumulates in more differentiated states (SPG.2-3 and SPG.4), suggesting an extended role beyond early spermatogonia. Our functional analysis revealed an association with lamellipodia, membrane protrusions that drive cell migration in both normal and pathological contexts. Given the established functions of PIWI proteins in stem cell maintenance and cellular migration ^40, 41^, this temporal shift suggests a more complex role for PIWIL4 during spermatogenesis than previously appreciated. These findings challenge assumptions based solely on transcriptomic data and demonstrate that protein-level measurements are essential for accurately defining cellular identity. In contrast, other genes such as the well-established spermatogonial marker RHOXF1 exhibit concordant RNA and protein expression patterns ^42^. RHOXF1 was identified as mainly expressed in SPG.0 with decreasing expression towards the SPC phenotype. This X-linked reproductive homeobox gene encodes a transcription factor with a critical role in regulating proliferation and differentiation, and mutations in this gene are associated with male infertility ^43^.

To further validate the mRNA-to-protein correlation of PIWIL4 and RHOXF1 observed by mIHC, we performed RNA-protein co-detection using RNAscope. By quantifying RNA punctate dots, we could confirm that *PIWIL4* and *UTF1* RNA were enriched in UTF1-protein high and PIWIL4-protein low SPGs. These data indicate that *PIWIL4* RNA is transcribed in earlier stages, thus revealing heterogeneous expression within the germ cell compartment. In contrast, *RHOXF1* RNA levels correlated with protein expression. These data highlight the dynamic transcriptional transitions associated with UTF1-, PIWIL4-, and RHOXF1-positive SPG populations.

Similar patterns were observed in spermatocytes, where both concordant and discordant RNA-protein relationships were detected. For example, the transcription factor RFX2 showed both mRNA and protein predominantly in SPC.3 cells, consistent with its known role during the pachytene stage and its involvement in acrosome assembly ^44^. Notably, the SPC.3 panel marker TCFL5 is also regarded as a pachytene protein, thereby increasing the reliability of our findings ^45^. Similarly, WNK1 is known to be expressed at the pachytene stage ^46^, further supporting the validity of our staining patterns. However, in contrast to its protein expression, WNK1 mRNA expression peaks were mainly detected in the SPC.1 state. WNK1 is essential for cell division ^47^, and disruption of WNK1 has been shown to induce premature protein synthesis of other mRNAs through upregulation of MTOR ^46^. Likewise, ACTRT3 displayed delayed protein expression relative to its transcript, consistent with previous reports of its localization in elongating spermatids ^48, 49^.

The coexistence of concordant and discordant expression patterns underscores the diversity of regulatory mechanisms governing gene expression during spermatogenesis. Periods of transcriptional inactivity, particularly during meiosis and spermiogenesis, necessitate reliance on stored mRNAs and tightly controlled translational activation. Well-characterized examples are Protamine 1 and 2 (PRM1 and PRM2), whose mRNA is transcribed in early spermatids but translated later due to regulatory elements within its 3′ UTR that maintain the transcript in a repressed state bound to inactive ribonucleoprotein particles. Premature translation disrupts spermatid differentiation, illustrating the importance of temporal control ^37, 50, 51^. Our data suggest that similar regulatory strategies may be more widespread than previously recognized, and the framework presented here provides a means to systematically identify such cases.

Beyond specific examples, our study demonstrates that integrating spatial proteomics with transcriptomics enables identification of regulatory programs that are not apparent from RNA data alone. This approach allows detection of delayed translation, stage-specific protein accumulation, and functional specialization across differentiation, thereby providing a more complete understanding of cellular identity. While our analysis focused on germ cells, spermatogenesis is supported by somatic niche cells, including Sertoli, Leydig, and peritubular (myoid) cells, which play essential roles in coordinating germ cell development and completing spermatogenesis ^52^. Future studies incorporating these cell types will be important for understanding cell-cell interactions and regulatory networks within the testicular microenvironment. In addition, expanding this framework to other tissues will further establish its utility for studying complex biological systems.

The methodological framework presented here is designed to be broadly accessible and scalable. By using fixed antibody panels combined with iterative staining of candidate proteins, the approach avoids the need for complex antibody conjugation and is compatible with standard histological workflows. Although each panel includes a limited number of markers, multiple panels can be combined to interrogate diverse cell types and states. Furthermore, automated image analysis enables quantitative and reproducible measurements, reducing subjectivity associated with manual interpretation.

Several limitations should be considered. The study is based on a limited number of adult human samples, which may not fully capture inter-individual variability or developmental differences relevant to infancy or pre-pubertal phases that could influence fertility. Limited clinical metadata may also introduce confounding factors related to medications or lifestyle. Furthermore, the analysis relies on static tissue sections, providing only a snapshot of protein expression rather than direct measurement of dynamic processes. Finally, although automated segmentation enables large-scale analysis, distinguishing closely packed or overlapping cells in transverse tissue sections remains challenging.

In summary, we present a scalable framework for spatial single-cell proteomics that enables quantitative mapping of protein expression across defined cellular states within intact tissue. By applying this approach to human spermatogenesis, we generate a high-resolution proteomic atlas and reveal widespread RNA-protein discordance, highlighting the importance of post-transcriptional regulation in defining cellular identity. These findings further show that multiple molecular layers are needed to accurately define cellular identity and function. More broadly, this work establishes a generalizable strategy for integrating transcriptomic and proteomic data to resolve cellular function and provides a foundation for future studies of tissue organization in health and disease.

## Methods

### Tissue preparation and ethical declaration

Histologically normal formalin-fixed and paraffin-embedded (FFPE) testis tissue samples from adult individuals were used for mIHC. The cohort comprised four individuals (#4485 Male 41 years, #4506 Male 32 years, #3627 Male 25 years, #4845 Male 29 years), of which three were included in each analysis. Testis tissues were obtained from the Department of Clinical Pathology at Uppsala University Hospital (Sweden) and collected through the Uppsala Biobank. Samples were anonymized in accordance with approvals from the Uppsala Ethical Review Board (ref nos. 2002-577, 2005-388, 2007-159), and informed consent was obtained from all subjects.

Tissue microarrays (TMAs) were generated by extracting 1 mm cores from FFPE blocks using a manual tissue arrayer (MTA-1, Beecher Instruments) and embedding them into recipient blocks. Each TMA contained six cores, comprising duplicate cores from three individuals. Publicly available IHC data from the HPA www.proteinatlas.org were generated using standard TMAs containing single cores from three adult testis samples. As samples across the HPA dataset originate from different individuals, not all IHC stainings derive from the same donors. FFPE TMA blocks were sectioned at 4 μm using a waterfall microtome (Microm H355S, ThermoFisher Scientific, Fremont, CA) and mounted on SuperFrost Plus slides (Thermo Fisher Scientific, Fremont, CA). Sections were dried overnight at room temperature (RT), baked at 50°C for 12-24 h, and stored at -20°C until use.

### Sample preparation and slide pretreatment

Slides were brought to RT, deparaffinized in xylene, and rehydrated through graded ethanol (99.9%, 96%, and 80%) to deionized water. Endogenous peroxidase activity was blocked using 0.3% hydrogen peroxide in 96% ethanol. Heat-induced epitope retrieval (HIER) was performed in a decloaking chamber (Biocare Medical, Walnut Creek, California, USA) at 125°C for 4 min in pH 6.0 Target Retrieval Solution (Agilent Technologies Inc., Santa Clara, California, USA). Slides were rapidly cooled to 90°C and then allowed to cool to RT prior to staining. Both chromogenic single-plex IHC (HPA standard) and mIHC were performed using a Lab Vision Autostainer 480S (Thermo Fisher Scientific, Fremont, CA). For mIHC, slides were subjected to a bleaching step after HIER using a solution of 0.2 M glycine and 1.5% hydrogen peroxide in TBS-Tween (Thermo Fisher Scientific, TA-999-TT). Slides were incubated in the bleaching solution in Falcon tubes under LED illumination with rotation for 1h at RT to reduce tissue autofluorescence. Slides were then maintained in TBS-Tween until staining. Chromogenic IHC using 3,3′-diaminobenzidine (DAB) was performed as previously described ^53^.

### Annotation of IHC images

Digital images of IHC-stained transverse sections of human testis from the HPA (v23.proteinatlas.org, Ensembl version 109) were used for extended annotation. In addition to antibody validation, inclusion criteria required at least moderate staining intensity in one or more testicular cell types. Manual annotation of the seminiferous epithelium was performed as previously described ^15^ and validated using a multi-class, multi-label hybrid feature-based deep learning model ^11^. For each antibody, staining intensity (strong, moderate, weak, negative), fraction of positive cells (>75%, 25–75%, 1–24%, <1%) and subcellular localization (nuclear, cytoplasmic, membranous) were recorded across eight testicular cell types. These included five germ cell types (spermatogonia, preleptotene spermatocytes, pachytene spermatocytes, round/early spermatids, and elongated/late spermatids), and three somatic cell types (Sertoli cells, Leydig cells, and peritubular cells). For the present study, analyses were restricted to germ cell types, and somatic cell annotations were excluded. Preleptotene and pachytene spermatocytes were combined into a single category for downstream analyses. All annotation data are provided in Supplementary Table 5.

### Single-cell transcriptomics

To identify and characterize human testicular germ cell populations, publicly available UMI count matrices were obtained from the Gene Expression Omnibus (GEO). Inclusion criteria were adult, healthy tissue samples generated using the 10X Genomics platform without prior cell sorting. Two datasets met these criteria: Guo et al. (2018) (^18^; GSE112013), comprising testis cells from three donors aged 17–25 years, and Nie et al. (2022) (^19^; GSE182786), including 12 donors aged 17–76 years. In the latter dataset, samples were grouped into young (17–22 years, n=4), older group 1 (62–66 years, n=5), and older group 2 (64–76 years, n=3). The third group was excluded due to reported spermatogenic impairment. Three UMI count matrices (one from GSE112013 and two from GSE182786) were imported into R and analyzed using Seurat v4 ^54^. Seurat objects were created for each dataset, requiring a minimum of 3 cells and 500 detected genes per cell. After standard quality control, cells with 500–5000 detected genes and <20% mitochondrial reads were retained. Each dataset was preprocessed independently following the Seurat RPCA integration workflow (https://satijalab.org/seurat/archive/v4.3/integration_rpca). Data were integrated using reciprocal PCA (RPCA), followed by UMAP dimensionality reduction and clustering. Clustering resolution values from 0.4 to 1.2 were evaluated, with a final resolution of 0.8 selected. Clusters were annotated based on established marker genes and assigned to major cell types (Supplementary Figure 1D). Germ cell populations - spermatogonia (SPG), spermatocytes (SPC), and spermatids (early and late SPT) - were extracted and re-analyzed separately, resulting in three independent Seurat objects (Supplementary Figure 2). Cell states within each group were assigned based on known marker expression. To ensure consistency with proteomic data, analyses were restricted to genes encoding proteins with detectable IHC staining (at least weak intensity) in germ cells. Cells lacking clear annotation were excluded. Cell counts for all clusters are provided in Supplementary Tables 1 and 2.

To identify potential markers for constructing the three mIHC panels, the Seurat function *FindAllMarkers* was used (Wilcoxon rank-sum test; min.pct = 0.20, logfc.threshold = 0.25, only.pos = TRUE) for each of the SPG, SPC, and SPT objects. Candidate markers were further prioritized based on fold change, defined as the ratio between the highest and second-highest mean expression across sub-clusters, with a cutoff of 1.5. These candidate lists were integrated with IHC data to select markers for mIHC panel design. Expression patterns across cell states were visualized using Seurat’s *DotPlot* function. To assess lineage progression and developmental trajectory of the identified germ cell populations, pseudotime analysis was performed using Monocle 3 (Supplementary Figures 1G and 2).

### Multiplex panel generation and staining

Panel marker antibodies were selected based on staining specificity, prioritizing proteins with germ cell-restricted expression and minimal signal in somatic or stromal compartments. Candidate markers were derived from differential expression analyses and cross-referenced with IHC data. To ensure clear discrimination between cell states, markers showing substantial overlap (i.e., a high proportion of double-positive germ cells) were replaced or reordered within panels to minimize co-expression.

Selected antibodies were initially validated in single-plex format to confirm compatibility with the OPAL detection system, preservation of staining patterns observed in chromogenic IHC, and sufficient signal-to-noise ratio. Antibodies failing these criteria were replaced with alternative clones targeting the same protein or with markers exhibiting similar cell type specificity. For multiplex panel design, a fixed OPAL fluorophore sequence was applied across all panels to streamline staining. Fluorophores were assigned as follows: cycle 1 (OPAL690), cycle 2 (OPAL620), cycle 4 (OPAL570), cycle 5 (OPAL480), and cycle 6 (OPAL-DIG+780). The third cycle was reserved for the candidate protein, which was iteratively exchanged for each staining. To optimize detection, low-abundance markers were preferentially assigned to brighter fluorophores, whereas highly expressed proteins were paired with dimmer fluorophores. Each antibody was further evaluated across all staining cycles to identify optimal positioning and to ensure consistent staining intensity and pattern.

Following bleaching and autofluorescence reduction, slides were subjected to mIHC staining at room temperature using the UltraVision LP HRP kit (Epredia). Each staining cycle included blocking, primary antibody incubation, horseradish peroxidase (HRP) polymer-conjugated secondary antibody incubation, OPAL fluorophore deposition (Akoya Biosciences, Marlborough, Massachusetts, USA), and antibody stripping in via HIER. After completion of all cycles, slides were incubated with OPAL 780-conjugated anti-DIG antibody and counterstained with 4′,6-diamidino-2-phenylindole (DAPI) (Invitrogen, D1306, Thermo Fisher Scientific). Slides were mounted using Prolong Glass Antifade mounting medium and cured overnight at RT prior to imaging. Imaging was performed using the PhenoImager system (Akoya Biosciences) according to the manufacturer’s instructions. Spectral unmixing and image export were conducted using inForm software (Akoya Biosciences) with a synthetic spectral library. For the SPG panel, a custom spectral library was generated due to the inclusion of OPAL650 in a subset of stainings. Detailed information on antibody panels, staining cycles, dilutions, and reagents is provided in Supplementary Table 6.

### Selection of candidate proteins for staining with mIHC panels

Candidate proteins were assigned primarily to a single panel based on cell type-specific staining patterns derived from the extended annotation of testis IHC images. Selection prioritized proteins with distinct and restricted expression within germ cell populations. Given the similarity in pretreatment and staining protocols between chromogenic IHC and mIHC, antibody dilutions for candidate proteins were initially guided by prior IHC optimization. In general, dilution factors were adjusted (∼1.5× relative to chromogenic IHC) to achieve optimal signal intensity in the multiplex setting. Staining performance was evaluated by comparison to corresponding chromogenic IHC patterns, ensuring consistency in localization and absence of non-specific signal. Slides were restained if panel marker expression was incomplete or deviated from established reference patterns based on manual inspection. Whenever possible, the same antibody clone was used for proteins analyzed across multiple panels. In cases where this was not feasible due to antibody availability or compatibility constraints, alternative antibodies (typically rabbit polyclonal antibodies) targeting the same protein were employed.

### Image Analysis

Fluorescence images were analyzed by processing each channel independently. Whole-slide images were batch-exported using Akoya Phenochart software (Akoya Biosciences) with either a synthetic or panel-specific spectral library, according to the manufacturer’s instructions. Each of the six TMA cores was exported as a separate TIFF file containing the following channels: DAPI, OPAL480, OPAL520, OPAL570, OPAL620, OPAL690, OPAL780, and autofluorescence. Nuclei were segmented from the DAPI channel using a pre-trained StarDist 2D model (nms_threshold = 0.7, probability_threshold = 0.486166) and used as proxies for individual cells. Mean fluorescence intensity values for all OPAL channels were then extracted for each segmented cell. Candidate protein expression (OPAL520 channel) was analyzed using a pixel classification approach implemented in Ilastik. A supervised classifier was trained on a subset of images (∼11% of the dataset) using three annotation classes (positive, negative, and background). The trained model was applied to all images to generate segmentation masks, which were used to assign candidate protein positivity at the single-cell level.

Panel marker channels were processed separately by converting each channel to grayscale and applying automated thresholding using Otsu or multi-Otsu segmentation. For multi-Otsu segmentation, the highest intensity class was defined as positive signal, while lower intensity classes were considered negative or background. Thresholding strategies were panel-specific: SPG panel images were processed using three-level multi-Otsu segmentation for all channels; SPC panel images were processed using standard Otsu thresholding; and SPT panel images were processed using three-level multi-Otsu segmentation for OPAL480, OPAL570, OPAL620, and OPAL690, and four-level multi-Otsu segmentation for OPAL780.

Segmentation performance was evaluated for each staining, and thresholds were adjusted when necessary to account for variability in signal intensity. Binary masks were generated for all channels, and manual inspection of a subset of images was performed to ensure accurate signal detection. Cells were phenotyped into defined germ cell states based on combinations of panel marker positivity derived from thresholded signals. Candidate protein expression was then overlaid onto all segmented cells, including both phenotyped and non-phenotyped populations. Each processed slide corresponded to one candidate protein within a specific panel, and the resulting single-cell data were compiled into data frames for downstream analyses. Mean fluorescence intensity values were calculated for each cell, and values corresponding to positive signal were retained for further analysis. Candidate protein expression values were normalized using min–max scaling to enable comparison across proteins within the same panel.

### Downstream descriptive analysis

Image analysis data from SPG, SPC, and SPT panels were processed in R (version 4.3.2). Data from each panel were organized into structured data frames, and metadata from the Laboratory Information Management System (LIMS) were used to exclude cells from low-quality tissue cores (e.g., lacking seminiferous tubules or sufficient marker representation). Cells were assigned phenotypes based on panel marker OPAL intensities according to predefined criteria (Supplementary Table 7), and cells without a defined phenotype were excluded from further analysis. For each cell type, mean candidate protein expression (OPAL520 intensity, scaled 0-1) and cell frequencies were calculated. Hierarchical clustering was performed on mean expression profiles for each panel, and resulting clusters were visualized as heatmaps using the pheatmap R package. Gene clusters were subsequently subjected to Gene Ontology (GO) enrichment analysis using the clusterProfiler package, considering Biological Process, Molecular Function, and Cellular Component ontologies. GO terms with an adjusted p-value < 0.05 were considered significant, and redundancy reduction was applied with a cutoff of 0.7. The top five enriched GO terms per cluster were visualized using lollipop plots, displaying the rich factor, adjusted p-value (color-coded), and gene count (point size).

### Transcript and protein expression dynamics

Correlation analysis was performed to assess the relationship between RNA and protein expression across germ cell differentiation states. scRNA-seq data were obtained from Seurat objects representing spermatogonia (SPG), spermatocytes (SPC), and spermatids (SPT), and protein expression data were derived from multiplex immunofluorescence (OPAL) measurements. Cells were grouped according to defined differentiation states. RNA expression values were normalized using min–max scaling (range 0–1), while protein expression values were used in their pre-scaled form. For each gene, summary statistics (mean, median, standard deviation, minimum, and maximum) were calculated across cell states.

To compare RNA and protein dynamics, expression values were z-score normalized across cell states for each gene. The normalized RNA and protein datasets were then merged by gene and cell state, and Pearson correlation coefficients were calculated to quantify concordance between RNA and protein expression profiles. Expression trends were visualized using line plots of mean expression across cell states with standard deviation ribbons. The relationship between correlation coefficients and peak RNA expression levels was visualized using scatter plots, and the distribution of correlation coefficients and peak RNA values was assessed using density plots (ggplot2).

### RNAscope and RNA-protein co-detection

Multiplex RNA in situ hybridization was performed on FFPE tissue sections using the RNAscope Multiplex Fluorescent Reagent Kit v2–Hs in combination with the RNAscope 4-Plex Ancillary Kit (Advanced Cell Diagnostics). A protease-free PretreatPro workflow was applied to preserve tissue morphology and enable RNA-protein co-detection. FFPE tissues were sectioned at 4-5 µm onto Superfrost Plus slides, air-dried overnight, and baked at 60 °C for 1 h. Slides were deparaffinized in xylene, rehydrated through graded ethanol, and rinsed in distilled water. Endogenous peroxidase activity was quenched using RNAscope Hydrogen Peroxide for 10 min at room temperature. Target retrieval was performed in 1× RNAscope Target Retrieval Buffer at 95–99 °C for 30–35 min (15 min for FFPE cell pellet controls), followed by immediate cooling. Sections were then treated with RNAscope Manual PretreatPro for 35 min at 40 °C in a humidified HybEZ oven.

#### Probe hybridization and signal detection

Following pretreatment, slides proceeded to probe hybridization. Three experimental configurations were used: (i) four-plex RNA detection, (ii) dual RNA detection combined with immunofluorescence, and (iii) dual RNA detection alone. The Channel 1 probe Hs-UTF1 (cat.no. 868061) was used ready-to-use, while Channel 2 probes (50x stock dilutions) were diluted into Channel 1 to generate probe mixes. The following probes were used: Hs-RHOXF1-C2 (cat.no. 1669551-C2) and Hs-PIWIL4-C2 (cat.no. 807661-C2). Probes were pre-warmed and hybridized at 40 °C in a humidified chamber. Signal amplification was performed using the RNAscope AMP1-AMP3 reagents, followed by sequential channel-specific HRP reactions. RNA targets were visualized using OPAL fluorophores (OPAL 520, 570, and 690), with HRP blocking steps between channels to prevent cross-reactivity. RNA integrity and assay performance were assessed for each experiment using multiplex positive and negative control probes. Positive controls targeted housekeeping genes (POLR2A, PPIB, and UBC) to represent low, medium, and high expression levels, while a bacterial dapB probe served as a negative control to monitor background signal (Supplementary Figure 6).

#### RNA-protein co-detection

For RNA-protein co-detection experiments, immunofluorescence staining was performed following RNAscope amplification. Sections were blocked for 1 h in 1X BSA blocking buffer (Thermo Fisher Scientific, cat. no 37520), followed by primary antibody incubation for 2 h with mouse anti-UTF1 (Merck, MAB4337, 1:500), rabbit anti-PIWIL4 (Atlas Antibodies, HPA036588, 1:100), and rabbit anti-RHOXF1 (Atlas Antibodies, HPA056506, 1:100). Secondary antibody incubation was performed for 45 min, and all signals were developed using OPAL 520. Nuclei were counterstained with DAPI (Invitrogen, D1306, Thermo Fisher Scientific). Slides were mounted with antifade medium and scanned using an Akoya PhenoImager (Akoya Biosciences) at 20x magnification. Multispectral images were unmixed using a synthetic library and exported as multipage TIFF files.

#### Image analysis

RNAscope experiments were performed on the same TMA used for mIHC. Image analysis was conducted using QuPath ^55^. At least 10 seminiferous tubules per staining were manually annotated, and cells were segmented using the InstanSeg segmentation tool. RNA puncta were detected using the “Subcellular Detections” function. RNA positivity thresholds were defined per slide based on visual inspection to account for staining variability. RNA expression was quantified as the number of puncta per cell. For RNA-protein co-detection analyses, cells were stratified into protein-high and protein-low populations based on manually defined intensity thresholds of the OPAL 520 channel, and RNA puncta counts were summarized separately for each group.

## Declarations

### Ethics approval and consent to participate

Testis tissues were obtained from the Clinical Pathology department, Uppsala University Hospital (Sweden), and collected within the Uppsala Biobank organization. Samples were anonymized as determined in the approval and advisory report from the Uppsala Ethical Review Board (ref nos. 2002-577, 2005-388, 2007-159). Informed consent was obtained from all subjects in the study.

### Consent for publication

Not applicable.

### Availability of data and materials

All data and images generated from this study are available in the Human Protein Atlas (https://www.proteinatlas.org/). All other data are available in the main text and the supplementary materials. All R codes and other source data are available upon request or at https://github.com/LindskogLab/A-spatiotemporal-atlas-of-human-spermatogenesis. The image analysis Python code is available on GitHub by accessing Jupyter notebooks and relevant src functions in this repository here: https://github.com/BIIFSweden/spermatogenesis-profiling (DOI:https://zenodo.org/doi/10.5281/zenodo.13788792).

### Authors’ contributions

Conceptualization: CL and MU; Data curation: FH and LM; Formal Analysis: FH, LM, GM, JG, BK, and RS; Funding acquisition: CL and MU; Investigation: FH, LM, JG, BK, and RS; Methodology: CL, FH, LM, GM, and JG; Project administration: JG and FH; Resources: LM, KvF, PA, GM, and MF; Soft-ware: LM, GM, FH; Supervision: CL, LM, CZ, and MU; Validation: FH, LM, and CL; Visualization: FH and LM; Writing – original draft: FH, LM, and CL; Writing – review & editing: FH, LM, CL, BK, RS, CZ, and MU.

## Acknowledgements

All members of the Human Protein Atlas are acknowledged for their efforts. We would also like to thank Uppsala University Hospital, and all patients with their families for donating tissue samples for research.

