## Supplementary figures and images for "A spatial single-cell type multiplex map of human spermatogenesis"

### Supplementary Figure 1

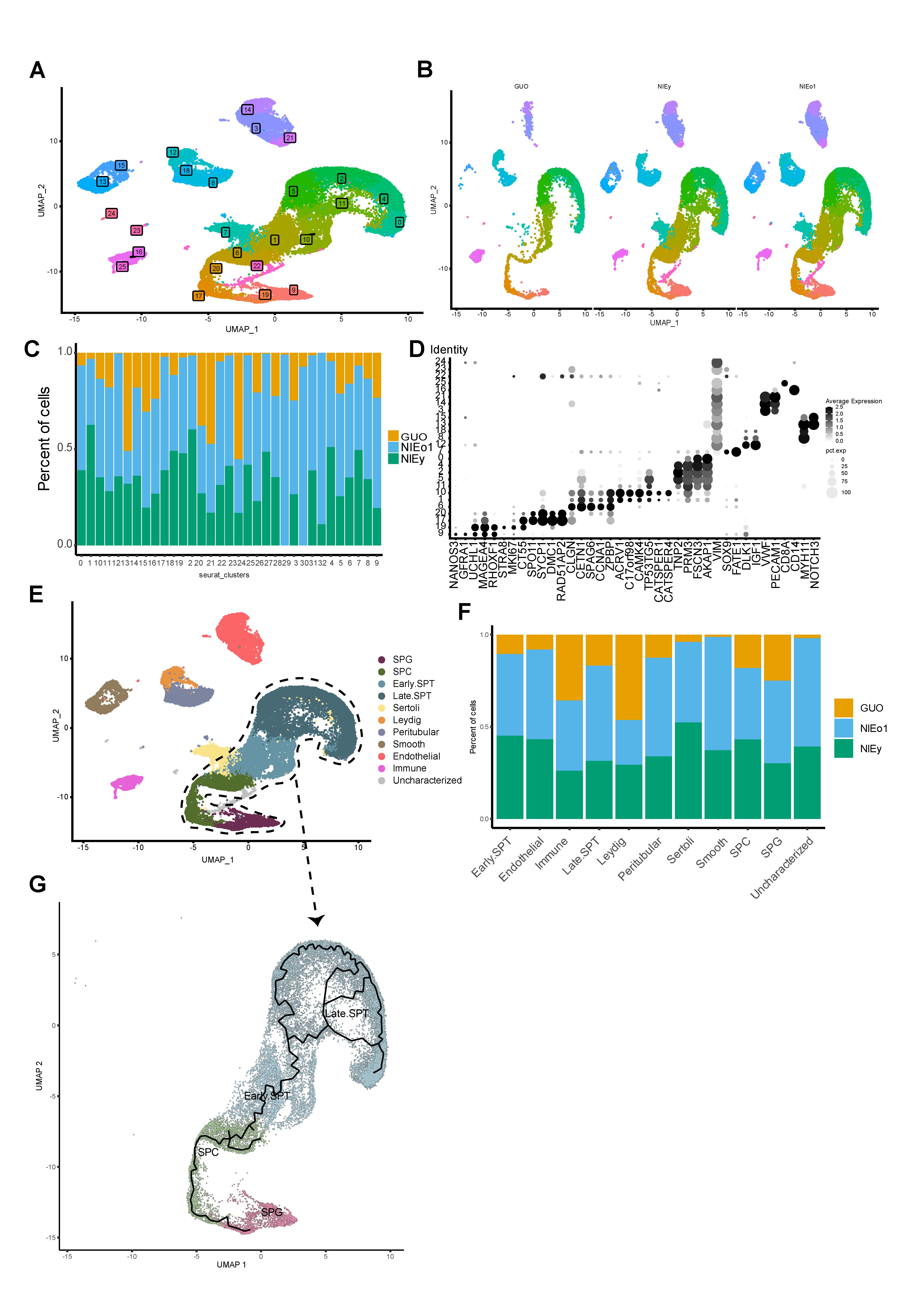

### Supplementary Figure 2

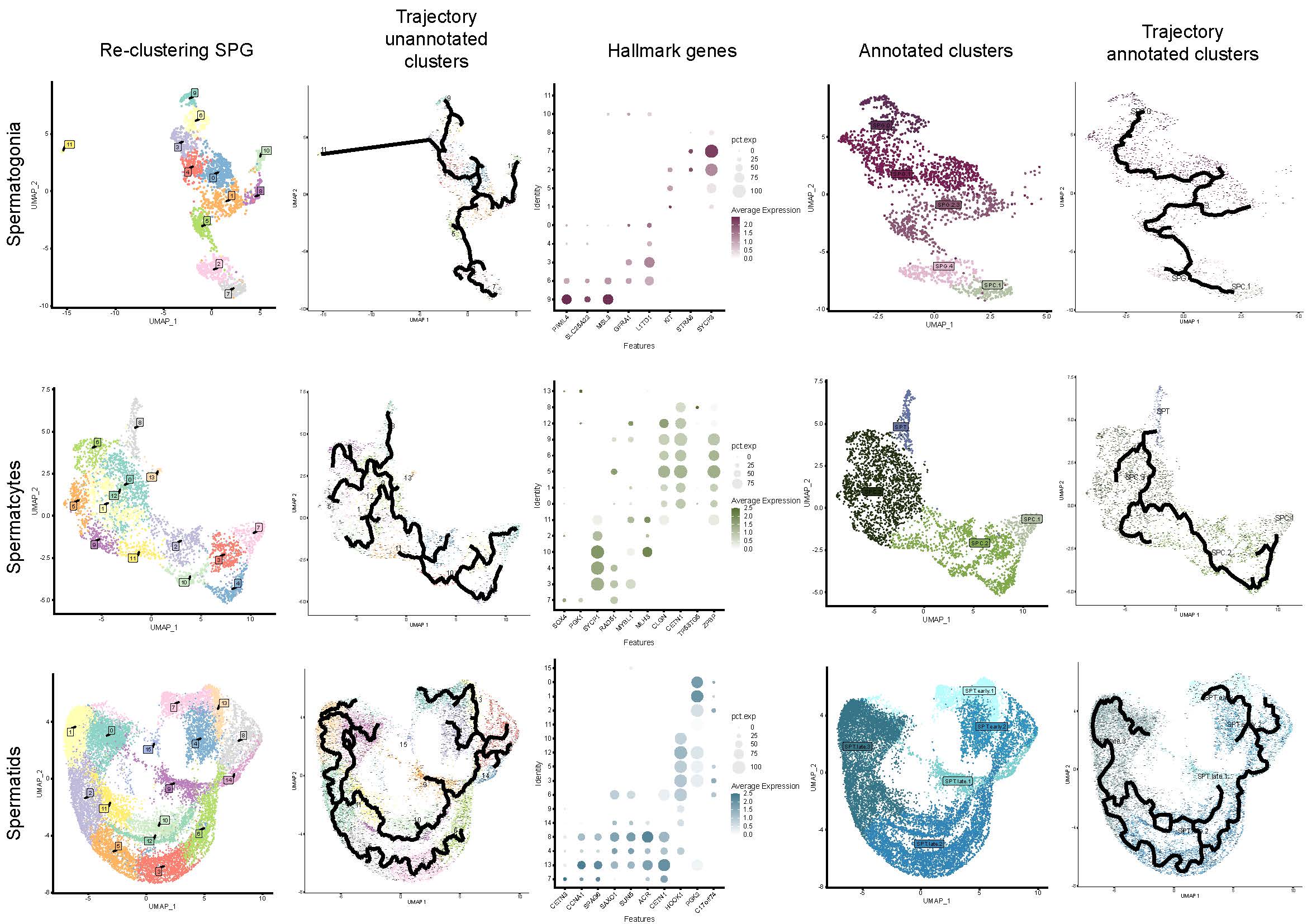

### Supplementary Figure 3

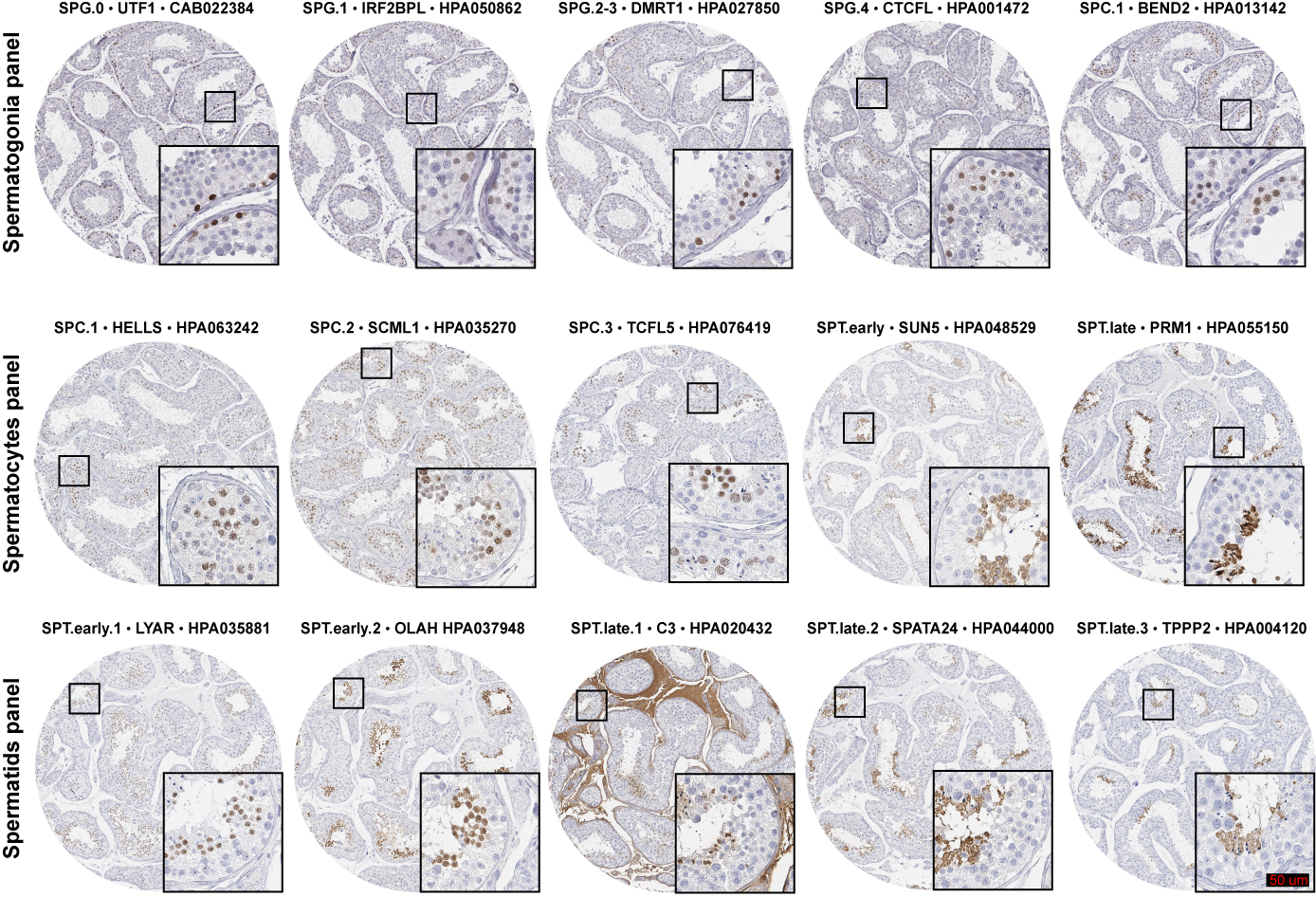

### Supplementary Figure 4

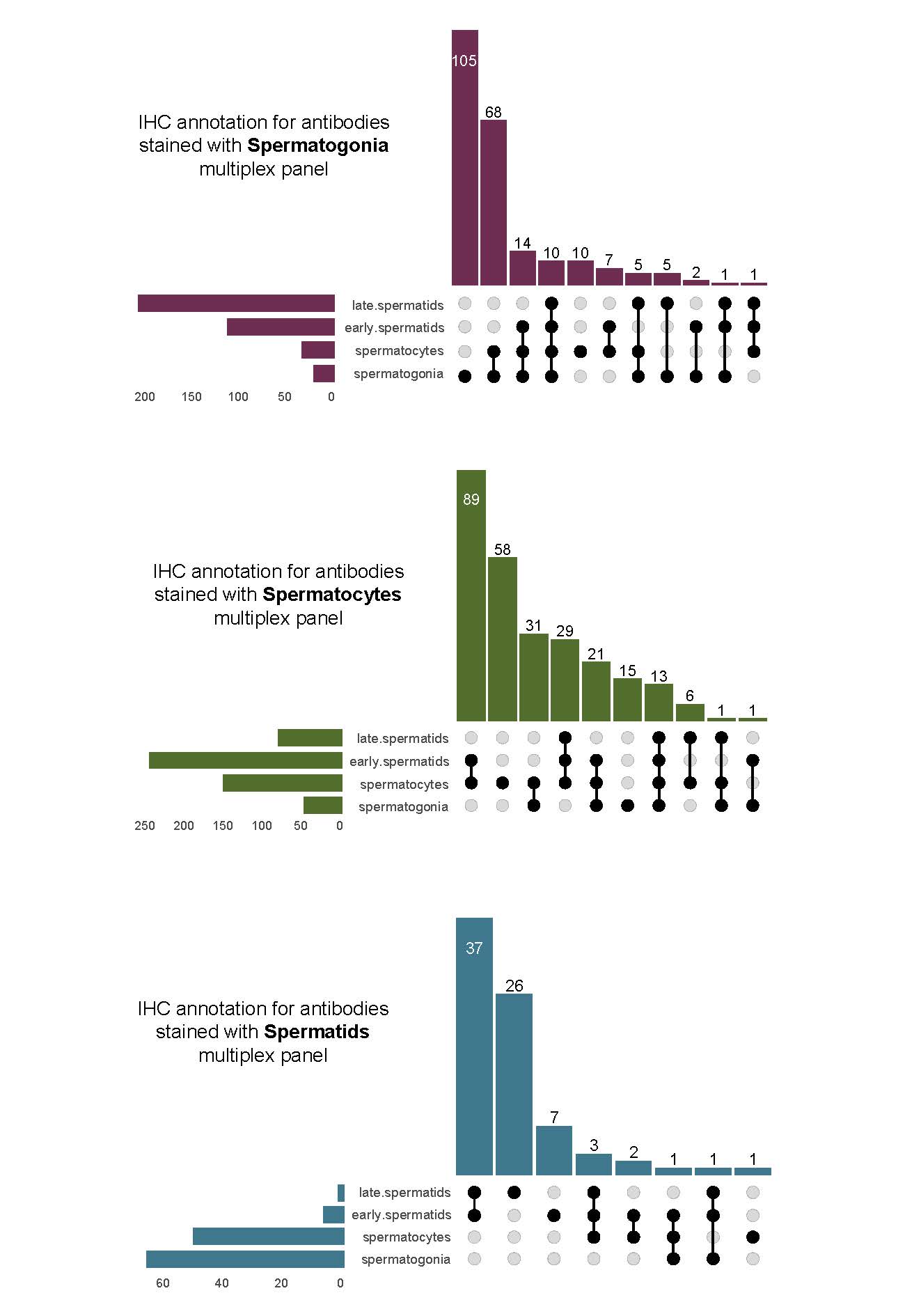

### Supplementary Figure 5

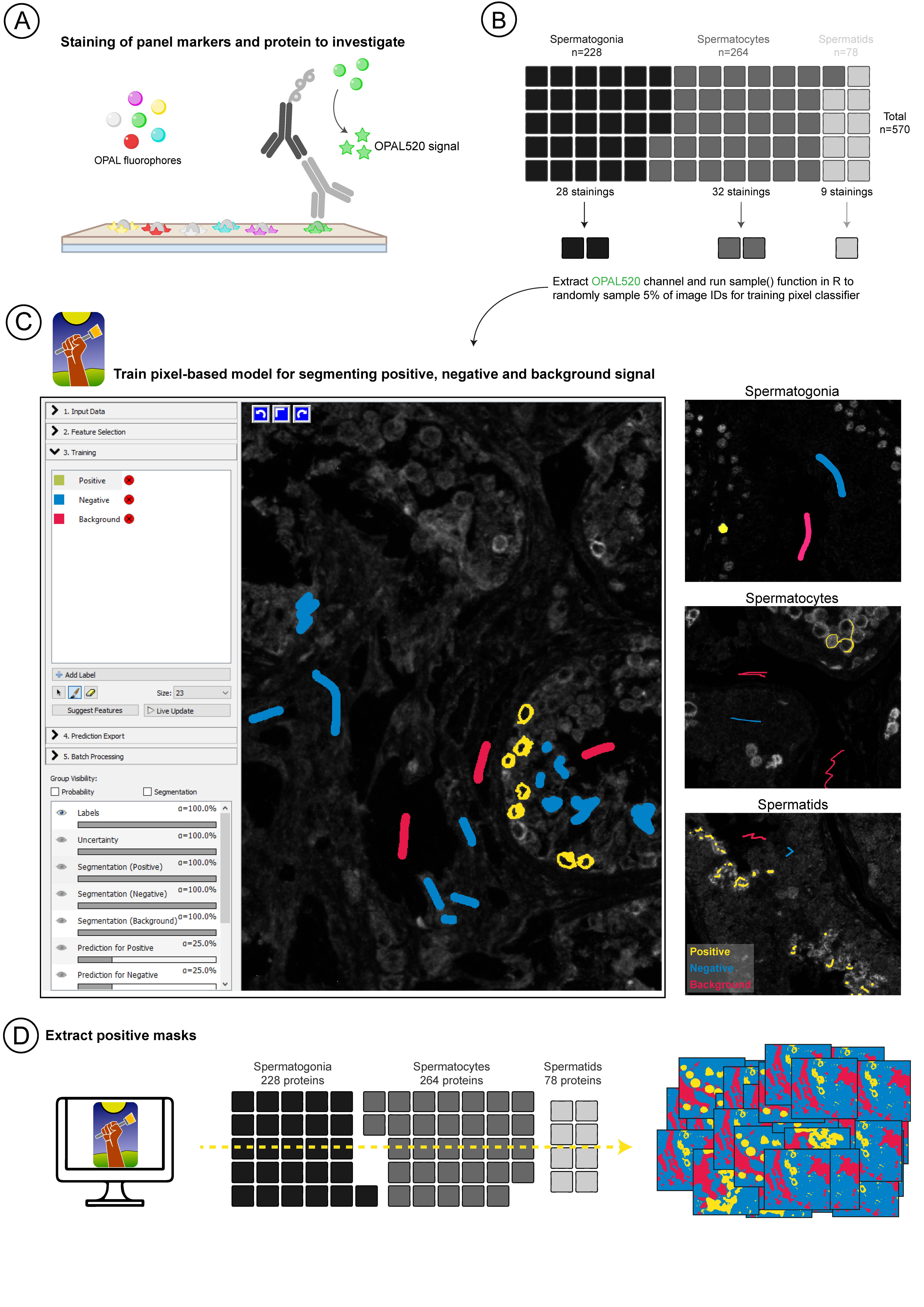

### Supplementary Figure 6

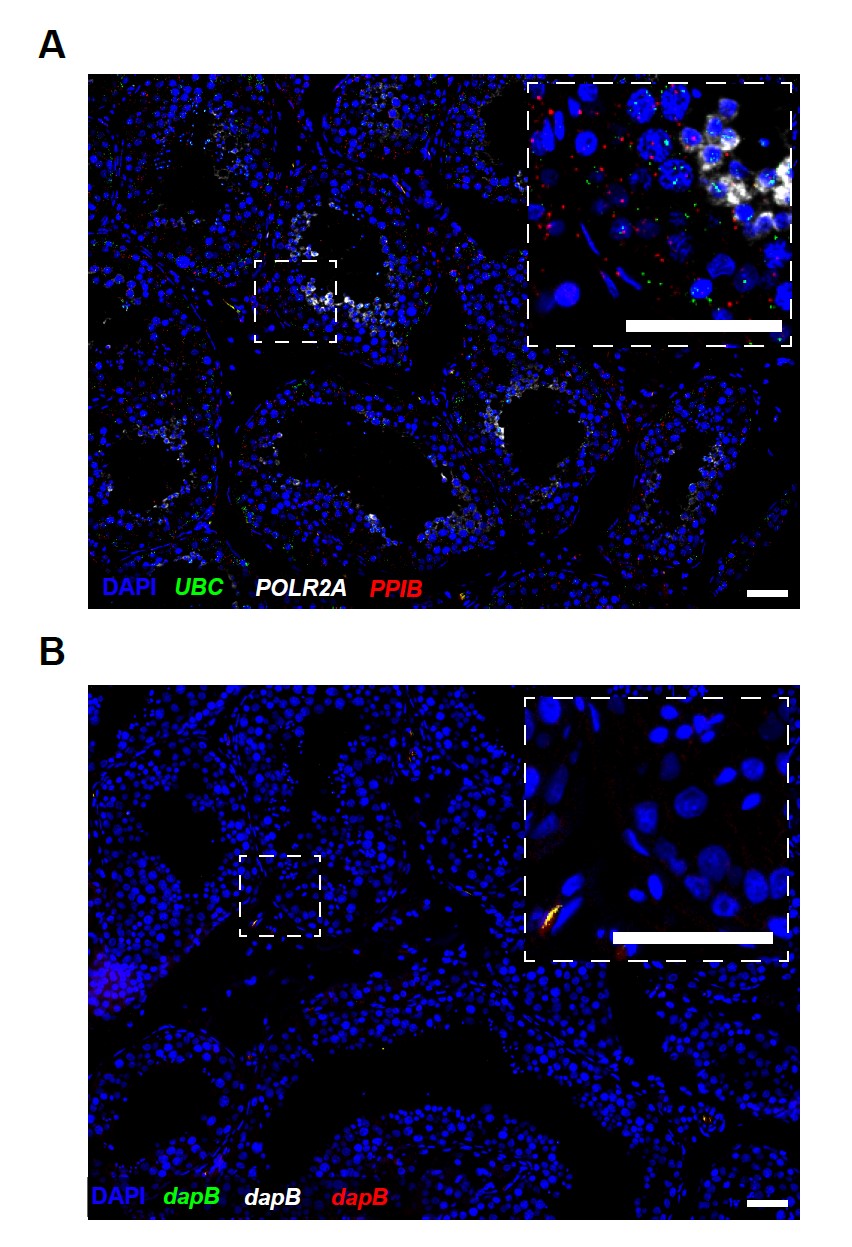
